# Data-driven modeling of core gene regulatory network underlying leukemogenesis in IDH mutant AML

**DOI:** 10.1101/2023.07.29.551111

**Authors:** Ataur Katebi, Xiaowen Chen, Sheng Li, Mingyang Lu

## Abstract

Acute myeloid leukemia (AML) is characterized by uncontrolled proliferation of poorly differentiated myeloid cells, with a heterogenous mutational landscape. Mutations in IDH1 and IDH2 are found in 20% of the AML cases. Although much effort has been made to identify genes associated with leukemogenesis, the regulatory mechanism of AML state transition is still not fully understood. To alleviate this issue, here we develop a new computational approach that integrates genomic data from diverse sources, including gene expression and ATAC-seq datasets, curated gene regulatory interaction databases, and mathematical modeling to establish models of context-specific core gene regulatory networks (GRNs) for a mechanistic understanding of tumorigenesis of AML with IDH mutations. The approach adopts a novel optimization procedure to identify the optimal network according to its accuracy in capturing gene expression states and its flexibility to allow sufficient control of state transitions. From GRN modeling, we identify key regulators associated with the function of IDH mutations, such as DNA methyltransferase DNMT1, and network destabilizers, such as E2F1. The constructed core regulatory network and outcomes of in-silico network perturbations are supported by survival data from AML patients. We expect that the combined bioinformatics and systems-biology modeling approach will be generally applicable to elucidate the gene regulation of disease progression.

**Significance:** A combined bioinformatics and systems-biology modeling approach is designed to model a transcriptional regulatory network for AML with IDH mutations. Network modeling identifies key regulators DNMT1 and E2F1, which is supported by patient survival data.

## Introduction

AML, the most common acute leukemia in adults, is characterized by uncontrolled proliferation of poorly differentiated and immature myeloid cells. Three classes of mutations have been observed in leukemic myeloid cells ^1^. Class I mutations are followed by class II mutations, contributing to about 80% of the AML cases. Class I mutations lead to the activation of receptor tyrosine kinases FLT3, KIT, and RAS signaling pathway, inducing cellular proliferation. Subsequent class II fusion mutations RUNX1/ETO, CBFB/MYH11, and PML/RARA affect transcription factors (TFs) RUNX1, CBFB, and PML and compromise normal differentiation. Class III mutations are found in genes encoding epigenetic modifiers such as DNMT3A, IDH1, IDH2, TET2, ASXL1, and EZH2, and can cause leukemia with worse patient outcome^1^. Specifically, mutations in IDH1 and IDH2, two genes encoding the cytoplasmic and mitochondrial forms of isocitrate dehydrogenase, respectively, are found in about 20% of AML cases^2^. These mutations contribute to a hypermethylated state in AML^3^. Moreover, IDH mutations and TET2 mutations are mutually exclusive^3, 4^ and IDH-mutant methylation and gene expression profiles are similar to those in TET2-mutant AML, suggesting a common pathogenic pathway^3^.

Although much effort has been made to elucidate the mutational landscape of AML and the linkage between these AML-associated mutations and disease severity, the gene regulatory mechanism of leukemogenesis is not yet fully understood. AML is a complex disease that arises from misregulation of gene regulatory network (GRN) driving normal cellular differentiation^5^. Therefore, mathematical modeling of the underlying GRN of AML and the effects of genetic perturbation can elucidate the gene regulation of the disease process and shed lights on new therapeutic strategies for AML. Some recent GRN modeling studies made efforts to elucidate AML gene regulation^6–12^. For example, Wooten et al. constructed a GRN of 106 nodes and 270 edges by composing interactions from different sources (e.g., SIGNOR) and performed Boolean modeling of the network to study drug response in class I FLT3 mutated AML^11^. Another recent Boolean network modeling study has refined a GRN model to recapitulate cellular state transitions during early hematopoiesis aging^13^. Despite the success of these modeling efforts, what is still missing is an approach that allows to systematically establish mechanistic models of GRN driving a specific subtype of AML. A promising solution to this question is to integrate top-down bioinformatics approach and bottom-up mathematical modeling for constructing GRNs of key transcription factors (TFs), referred as core GRNs^14^. A recently developed method, named NetAct^62^, has adopted this approach for modeling core GRNs driving cellular state transitions using gene expression data of multiple states and literature-based TF-target databases. Further generalization of this approach to integrate context-specific transcriptomics and epigenomics datasets and to enable GRN model selections based on network dynamics would allow to improve its capability for generating high-quality context-specific network models.

Here, we developed a new data-driven approach to inferring and modeling GRN regulating leukemogenesis in IDH1/2 mutated AML by integrating top-down bioinformatics approach and bottom-up mathematical modeling^14^. We first integrated data from diverse sources, including a microarray gene expression dataset, an ATAC-seq (Assay for Transposase-Accessible Chromatin using sequencing) data set for genome-wide chromatin accessibility, and literature-based TF to target gene relationship databases, to infer putative GRNs. For each GRN, we then applied a mathematical modeling method named *ra*ndom *ci*rcuit *pe*rturbation (RACIPE)^15–18^ to simulate the expression profiles of network genes for an ensemble of models with diverse kinetic parameters. The modeling approach has been streamlined to allow for a high-throughput application to many GRN topologies derived from the bioinformatics methods. We then identify the optimal GRN model where simulated gene expression data best match the experimental data, and meanwhile the GRN is sufficiently flexible to allow control of state transitions. From the established optimal GRN, we performed network perturbation modeling to identify key regulators associated with the mechanistic function of IDH mutations, such as DNMT1, and network destabilizers, such as E2F1, which are supported by patient survival data. Our modeling analysis further identifies the presence and coupling of key biological pathways, such as cell cycle, AMPK, and p53 pathways. In short, the combined bioinformatics and systems biology modeling approach has allowed to uncover key factors underlying leukemogenesis.

## Materials and Methods

### Integrative network modeling framework

We designed a new computational network modeling framework that integrates bioinformatics methods with mathematical modeling to infer context specific gene regulatory network (GRN). The framework consists of the following steps, as illustrated in **Fig. 1**. First, key TFs are identified by applying three distinct network construction methods, namely VIPER^19^, RI^20^, and NetAct^21^ (details in **Supplementary Note 1**).

**Figure 1.**
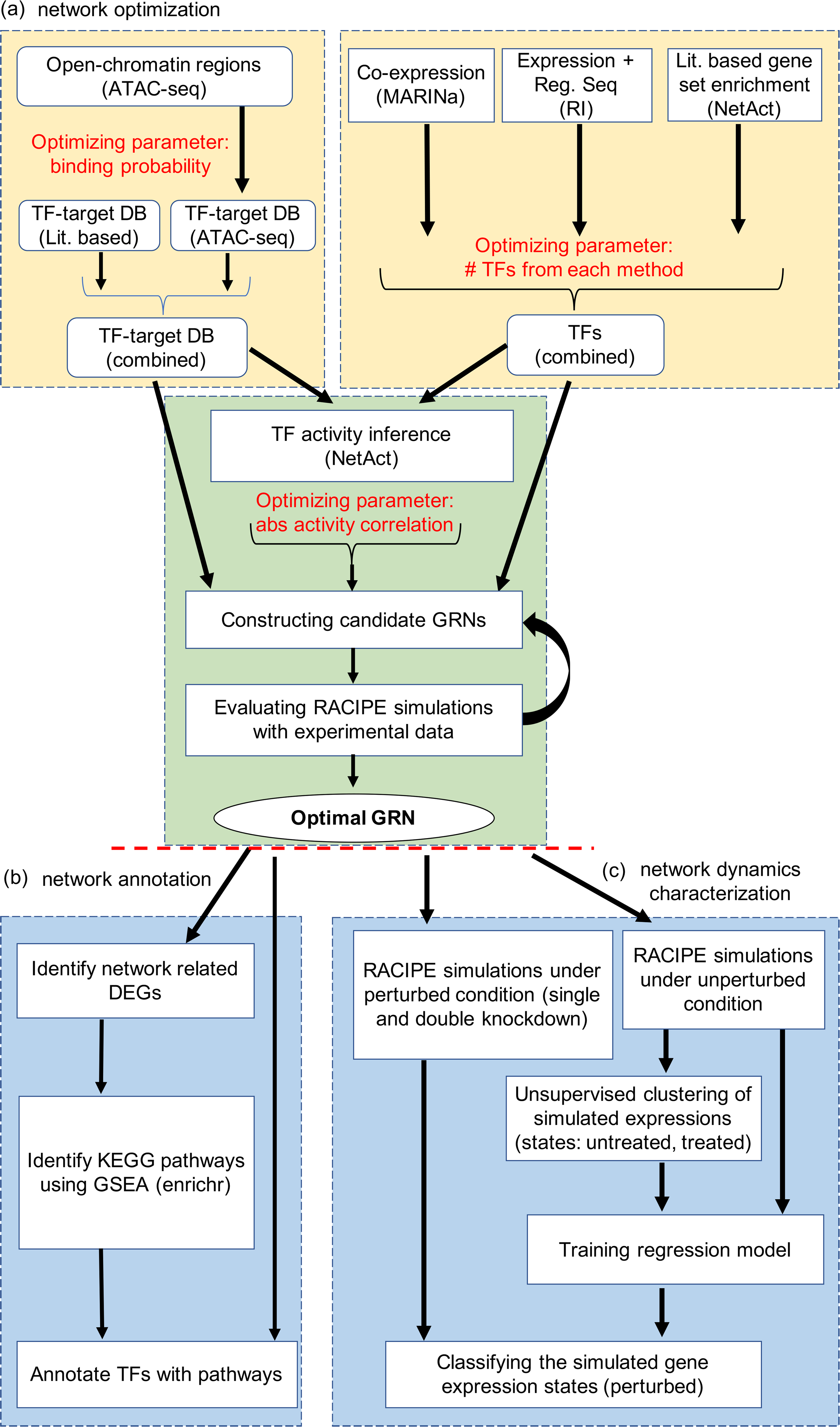
Illustration of the computational framework for gene regulatory network modeling. **(a)** Procedures for gene regulatory network (GRN) optimization. The top left block shows the steps to construct TF-target databases (DB) using a literature-based TF-target DB and the TF-target relationships inferred from ATAC-seq data. The top right block shows the approach of TF inference using three distinct computational methods: VIPER, RI regression, and NetAct. The bottom block shows the steps to construct GRN candidates using the TF-target databases and TF activities. Many candidate GRNs are constructed by varying three adjustable hyperparameters, as highlighted in red color. Network optimization is then applied to identify the optimal GRN that best captures experimental gene expression states according to GRN simulations by RACIPE. **(b) Network modeling analysis.** Using the GRN-related differentially expressed genes (DEGs), we identify enriched KEGG biological pathways and the best representative pathways associated with each network TF (left block). In silico network perturbation analysis can be further performed to identify key regulators of the network driving state transitions (right block).

Second, a context-specific TF-target database is constructed by combining curated TF-target databases and TF-target gene relationship derived from ATAC-seq data (details in **Supplementary Note 2**). Third, the activity of each key TF is inferred by NetAct using the expression of their corresponding target genes. Fourth, a GRN consisting of the key TFs is constructed, where a regulatory link between two TFs is determined by the correlation of the activities of the TFs. We sampled three network construction parameters, namely ATAC-seq TF-binding probability cutoff, number of TFs taken from each TF selection method, and correlation cutoff of TF activities (**Fig. 1a**), which generated 532 candidate GRNs (details in **Supplementary Notes 3 and 4**). Subsequently, we applied the mathematical modeling method RACIPE^18^ to each GRN to evaluate how well the GRN steady states capture the TF activity profiles from both the normal controls and the AML patients. We used enrichr ^22^ to find the significantly enriched biological pathways in the differentially expressed genes (with adjusted p-value <= 1.60e-10) and annotated the TFs with the most representative pathways (Fig. 1b). Finally, network simulations and gene perturbation analyses were performed on the optimized GRN to predict the key regulators, which can be potential therapeutic targets of AML (**Fig. 1c**). More details on network annotation and network dynamics characterization can be found in **Supplementary Notes 5 and 6.**

### Gene expression data

We used a previously published microarray gene expression data for the primary AML patients (n = 119) and a control group from normal bone marrow CD34+ hematopoietic stem and progenitor cell (HSPC) specimens (n=11), which was profiled using Affymetrix Human Genome U133 Plus 2.0 GeneChips (Gene Expression Omnibus (GEO) accession number GSE6891)^23, 24^. In this study, raw data were reprocessed using the HGU133plus2.0 BrainArray annotation version 17.0.0. Gene expression levels were transformed to log2 values. Network modeling analyses were applied to the data for *IDH-*mutant AML patients (n=9, IDH1/IDH2 mutation and without DNMT3A mutation) and the normal controls to identify context-specific TFs.

### ATAC-seq data

We utilized ATAC-seq data to identify open chromatin regions within the promoter region, enabling the identification of context-specific TF-target relationships. The ATAC-seq datasets for leukemia stem cells from seven AML patients were obtained (GEO with accession number GSE74912)^25^. Sequencing data were pre-processed by the interactive-ATAC (I-ATAC) pipeline^26^. Briefly, we used *Trimmomatic*^27^ to identify and trim adapter sequences and low quality nucleotide sequences from the raw ATAC-seq read. Trimmed reads of each sample were mapped to the human reference genome GRNh37/hg19 by *BWA*^28^. *Picard* (https://broadinstitute.github.io/picard/) was used to filter PCR duplicated reads and calculate inset size. Next, I-ATAC adjusted sequencing as described by pipeline^26^ and the outcome was converted into the BED format to identify genomic regions enriched in the putative open chromatin sites (peaks) by *MACS*^29^. Finally, the ATAC peaks presented in all the seven AML patient datasets were used for TF binding site prediction.

### Survival analysis

In order to determine whether important TFs identified by our algorithm are associated with complete remission in AML, we used gene expression and clinical information for 119 primary AML patients^24^. First, a univariate Cox regression analysis was performed to evaluate the association between expression levels of genes and event-free survival of AML patients (event denotes failure to achieve complete remission). Then, we calculated a risk score for each sample which was defined as a linear combination of expression values of genes in one signature set weighted by their estimated Cox model regression coefficients. If the risk score for one sample was larger than the median risk scores, then it was classified into a high-risk group, otherwise into a low-risk group. Finally, Kaplan-Meier survival estimation and log-rank test were applied to evaluate the differences in patients’ survival time between the high-risk group and the low-risk group.

## Results

### Mathematical modeling identifies the optimal GRN

We inferred key TFs by applying three distinct methods, VIPER, RI, and NetAct, to analyze the microarray gene expression profiles from a cohort of nine AML patients with *IDH* mutations and eleven normal controls. First, we obtained a ranked TF list by applying VIPER, which assesses TF activity by combining transcriptional activation of its activated and repressed targets and its biological relevance by the targets overlapping with phenotype-specific programs (**Fig. 2a**). We obtained the second TF list by applying the regulator inference (RI), a lasso regression-based method, to the gene expression data and the TF motif binding sites from the ATAC-seq data. This method assigns importance score to each TF (**Fig. 2b**). We then obtained the third TF list by applying NetAct, which identifies the enriched TFs by performing gene set enrichment analysis (GSEA, with slight adjustments^21^) using a curated TF-target database on the differentially expressed genes between the normal controls and the AML patients with IDH mutations (**Fig. 2c**). These three methods (VIPER, RI, and NetAct) utilize different input datasets and capture different aspects of the underlying regulatory mechanism (see **Supplementary Note 1**).

**Figure 2.**
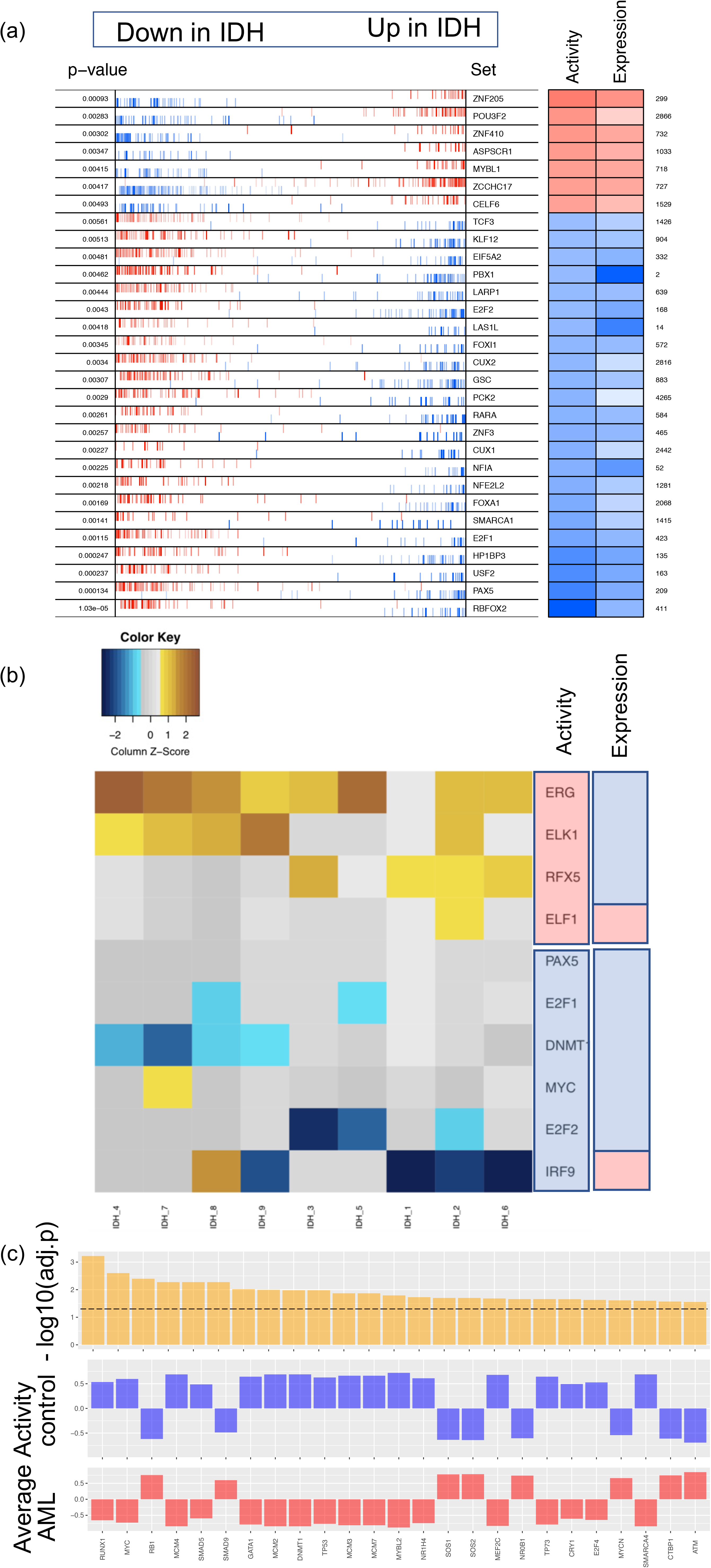
Enriched transcription factors identified from three different inference methods. **(a) VIPER**: Left side of the plot shows the distribution of the positively (red) and negatively (blue) correlated targets for each selected TF on the gene list ranked from the one most down-regulated to the one most upregulated in AML samples with *IDH* mutations compared with the samples of normal control. The two-column heatmap on the right side shows the inferred differential activity (first column labeled as Activity) and differential expression (second column labeled as Expression). **(b) RI**: The heatmap shows the AML sample-specific lasso model coefficients for each selected TF. In the annotation bars from the right side, the first column shows the activity of the TFs, and the second column shows the gene expression of the TFs (pink denotes upregulation, and blue denotes downregulation). **(c) NetAct:** 1^st^ row shows - log10(adjusted p-value) of the top 25 TFs ordered based on adjusted p-values; 2^nd^ row shows the average activities of normal control samples; 3^rd^ row shows the average activities of IDH samples. A horizontal dotted line represents adjusted p-value = 0.05.

From the inferred TFs by each method, we obtained many candidate GRNs of different sizes as follows. First, we constructed a combined TF-target gene-set database, which included literature-based TF-target gene sets and the TF-target gene relationships obtained from the ATAC-seq data at different TF-target gene binding probability threshold (see **Supplementary Note 2**). Next, we employed NetAct to calculate the activities of the selected TFs using the expression of their corresponding target genes, as defined by the combined TF-target database. Then, the calculated TF activities were used to infer candidate GRNs. The rationale behind using the TF activity, but not the expression, is that aberrant TF behavior in the disease state may not get manifested in the differential gene expression of the TF, rather in the coordinated activation of the target genes^30, 31^. We obtained 532 candidate GRNs by varying the hyperparameters – namely, the number of TFs selected from each method (VIPER, RI, NetAct), the ATAC-seq TF-target gene binding probability, and the TF activity correlation cutoff. Lastly, we used mathematical modeling to identify the optimal GRN whose simulated gene expression profiles best match the experimental data. To identify the optimal GRN, we applied RACIPE to each candidate GRN to generate an ensemble of 10,000 ordinary differentiation equation (ODE) models with randomly generated kinetic parameters (see **Supplementary Note 3**). Compared with the conventional modeling approaches where a set of kinetic parameters needs to be specified, RACIPE uses the topology of a GRN as the only input for modeling and identifies the network states from the gene expression clusters observed in the gene expression profiles from the ensemble of models^15–17^.

Using the simulated gene expression profiles from the candidate GRNs, we then ranked each GRN with two metrics, namely accuracy and flexibility. Here, the accuracy of a GRN is calculated as the proportion of the RACIPE-simulated gene expression profiles that match the experimental TF activity profiles ^32^ (**Fig. 3a**). This determines how well the simulation of a candidate GRN reconstructs the experimental data. We also defined flexibility^33^, which measures the average deviation of the proportional of models in the two states (i.e., normal and AML states) between the perturbed and unperturbed conditions over all gene knockdown simulations. A network with fewer connections will have higher flexibility than a dense network (**Fig. 3b**). See **Supplementary Note 4** for the calculation details. The distributions of accuracy and flexibility across the aforementioned three network construction parameters are shown in **Fig. 3c**. The optimal GRN is expected to exhibit high accuracy to capture the gene expression states and high flexibility to allow flexible control of state transitions. Therefore, we order the candidate GRNs based on both metrics, first by accuracy and then by flexibility, to obtain a combined ranking from both the metrics (see **Methods**). **Fig. 4a** shows the scatter plot of accuracy ranking versus flexibility ranking, where the optimal network is highlighted in red. Additionally, the optimal GRN stays as the top network over repeated simulations and re-ranking and is significantly different from the second-best networks (t-test, p-value < 0.05, **Fig. 4b**), suggesting convergence of the network optimization. The optimal GRN consists of 29 TFs and 102 regulatory interactions, of which 53 are excitatory and 49 are inhibitory (**Fig. 4c**). In the optimal GRN, 28% of the interactions are derived from the ATAC-seq data (28 out of 102 interactions).

**Figure 3.**
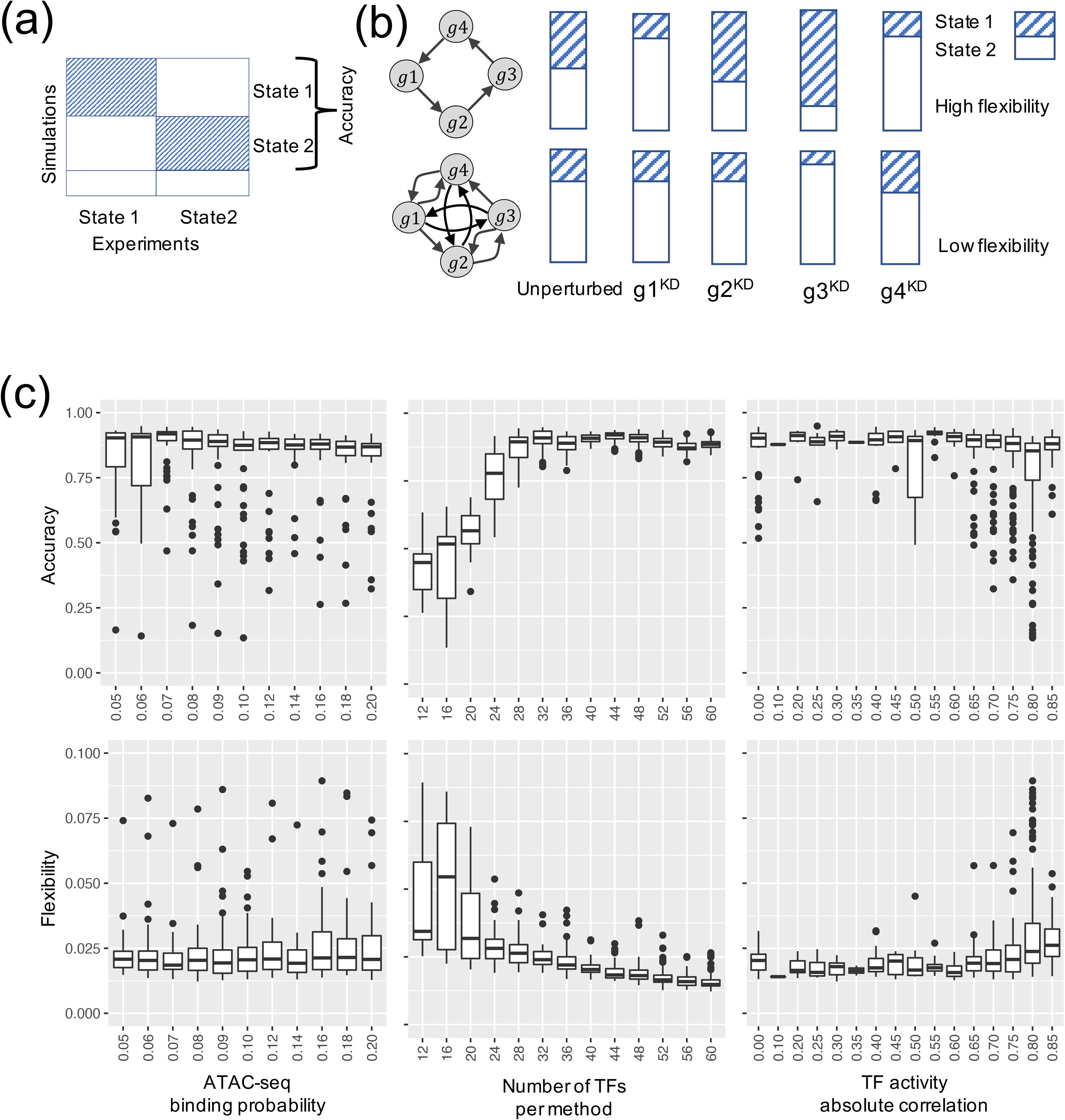
Network optimization by the accuracy and flexibility scores. **(a)** Schematic for the definition of accuracy. Accuracy is the fraction of the RACIPE models that can be assigned to any of the two clusters (states 1 and 2) of gene expressions. **(b)** Schematic of the definition of flexibility. Flexibility is measured by the average deviation of the proportional of models in the two states between the perturbed and unperturbed conditions over all gene knockdown simulations. The circuit with larger average deviation (top) is more flexible than the other circuit (bottom). **(c)** Distribution of accuracy (top panel) and flexibility (bottom panel) of candidate GRNs with respect to the optimization hyperparameters: TF-binding probability (leftmost panels), number of TFs per method (middle panels), and the correlation cutoff (absolute value) of TF activity (rightmost panels).

**Figure 4.**
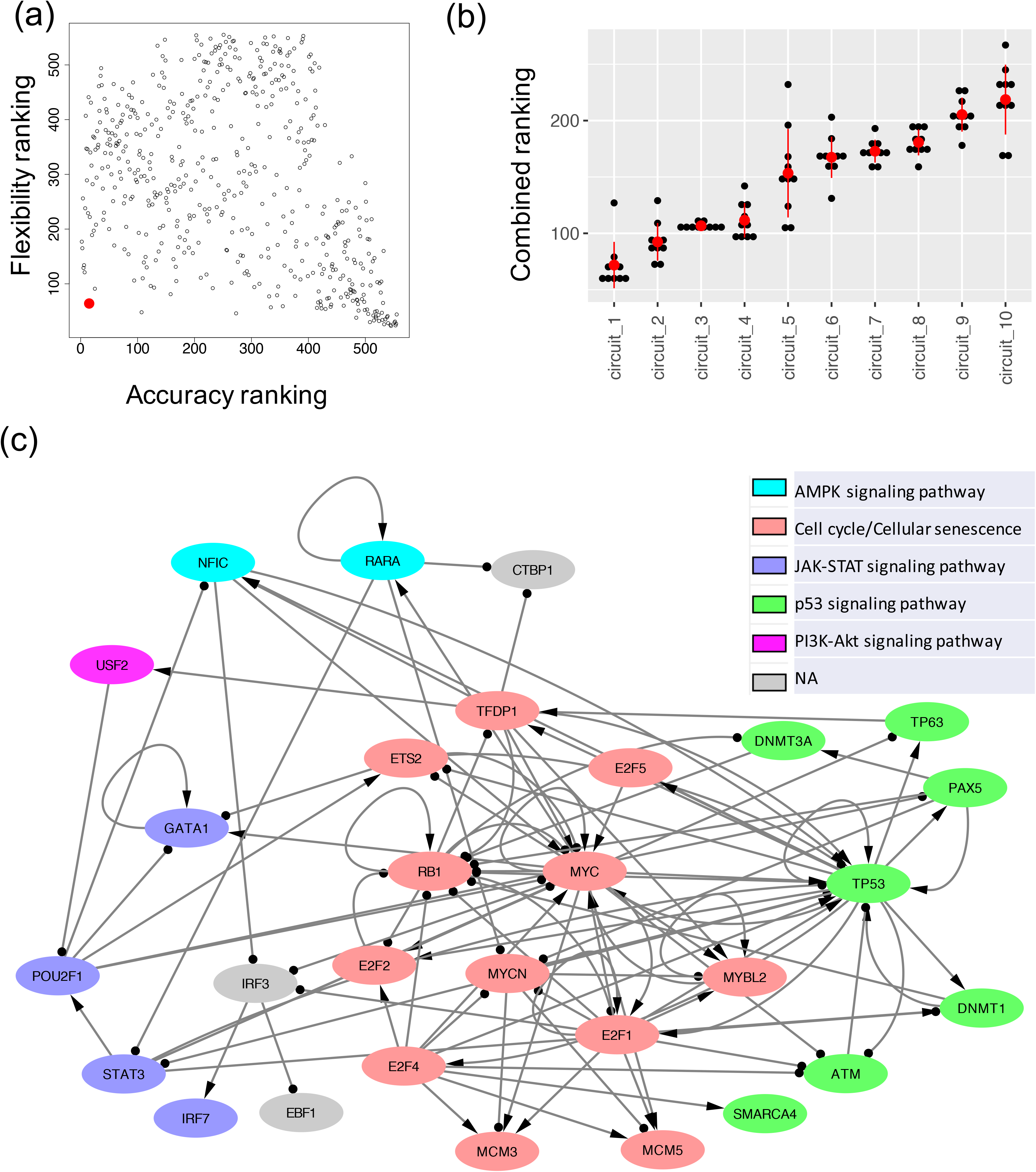
The optimal GRN of leukemogenesis in IDH mutant AML. **(a)** A scatter plot showing the accuracy ranking (x-axis) and flexibility ranking (y-axis) for a total of 532 GRN candidates. The optimal GRN is marked with the red enlarged dot. **(b)** Convergence of the top ranked networks. Distribution of the combined scores (sum of two rankings, one based on accuracy and the other based on flexibility) for the top ten GRNs obtained from the ten repeats of 10000 RACIPE simulations for each circuit. The red dot and the vertical bar are mean and standard deviation of the distribution for each circuit. A two-sided t-test shows that the scores for the top ranked GRN is significantly different from those of the other GRNs. **(c)** The diagram of the optimal AML GRN of enriched TFs. Transcriptional activation is annotated by a line with arrowhead; transcriptional inhibition is annotated by a line with circle head. The colors of the gene nodes represent the most representative KEGG biological pathways. The coupling of biological pathways is shown in **Fig. S2**.

### Simulations of the optimal GRN agrees well with the experimental data

We used NetAct to calculate the activities of the 29 TFs in the optimal GRN for the normal controls and the *IDH*-mutant AML patients. From the profiles of the activities and the expressions of the TFs that are included on the GRN (**Fig. 5a**), it is evident that the TF activity profiles can distinguish the normal controls and the AML patients well. Furthermore, RACIPE simulation of the optimal GRN shows high agreement with the experimental data. Here, to perform the similarity analysis, we generated 10000 gene expression profiles from RACIPE simulations of this network and then mapped the models to the TF activity profiles of either the normal controls or the AML patients (see **Supplementary Note 4** for profile mapping details). There is a subset of the RACIPE models (**Fig. 5b**, cluster with black marker at the top-right) that could not be mapped to any of the two groups, normal controls and AML patients. The lower the proportion of these unmapped models, the better the GRN captures the gene expression states of normal and cancer conditions. The accuracy of the optimal GRN, measured as the percent of models that conform with the data, is 0.93, where the proportions of the models that match the normal and cancer conditions are 0.24 and 0.69, respectively (**Fig. 5c**).

**Figure 5.**
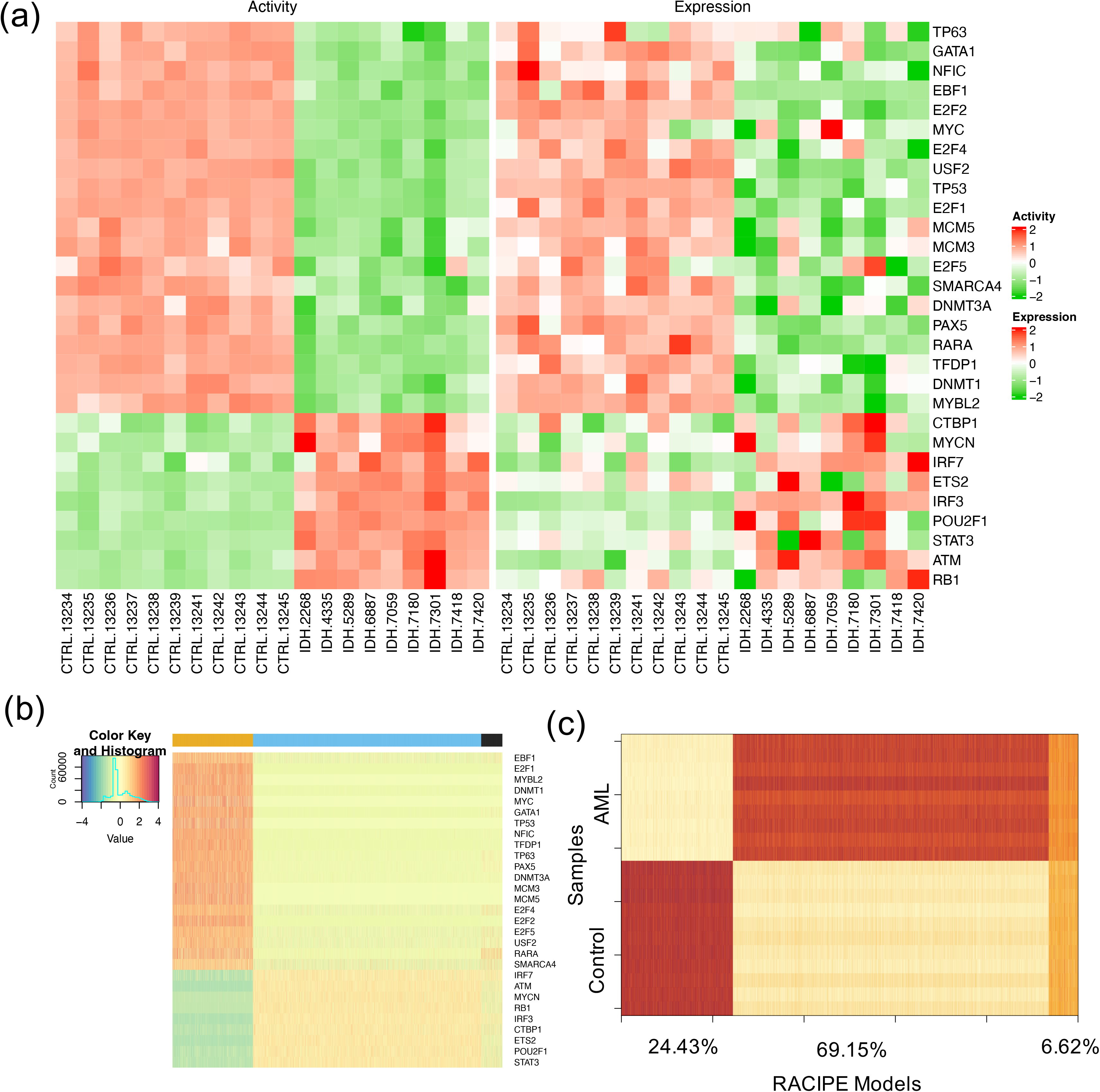
Simulation and characterization of the AML GRN. **(a)** Heatmaps showing the profiles of TF activities (left panel) and experimental gene expressions (right panel) for the TFs in the optimal GRN. For row clustering, Euclidean distance and complete linkage method were applied. The columns show sample names for the control and the AML samples. **(b)** Heatmap of the RACIPE simulated gene expression profiles for the optimal GRN. Hierarchical clustering analyses were performed with the distance of one minus Spearman correlation and complete linkage. **(c)** Spearman correlations between the TF activities across samples (11 normal controls and 9 AML patients) along y-axis and the RACIPE simulated gene expressions along x-axis. The percentages (24.43%, 69.15%, and 6.62%) along the x-axis are the percent of the RACIPE models that are mapped to control group, treatment group, and neither of the two groups, respectively.

### GRN modeling elucidates the drivers of leukemogenesis in IDH1/2 mutant AML

The optimal GRN associated with leukemogenesis in IDH1/2 mutant AML reveals the importance of DNMT1 as a key TF. Studies have shown that IDH1/2 mutations and TET2 mutations are mutually exclusive, resulting in an overlapping hypermethylation signature^3^. The oncometabolite 2-HG, produced by mutant IDH1/2, disrupts TET2 function and promotes oncogenesis^34^. Additionally, IDH1/2 mutations activate HDAC1/2, inhibiting the formation of the DNMT1 and TET2 complex, leading to the degradation of DNMT1 and TET2^35^. This impairment of the DNMT1 and TET2 complex formation contributes to abnormal DNA methylation in IDH-mutated AML. Moreover, the optimal GRN involves crucial cell cycle and DNA-damage-repair genes, such as RB1, E2F1/2, TP53, and MYC, and several stem cell pluripotency factors GATA1^36^, POU2F1, and MYCN^37^. The over expressions of these genes suggest that the AML cells attain stem cell like phenotype with a much-restricted cell cycle, which may induce drug resistance to these AML cells^38, 39^. These TFs can also facilitate the coupling of multiple pathways to carry out the required complex biological functions.

### GRN modeling identifies the presence and coupling of key biological pathways

Furthermore, we identified six key KEGG pathways^40^ involving the TFs in the optimal GRN by performing GSEA using the TFs and their target genes (details in **Supplementary Note 5**). These enriched pathways include two regulatory pathways (cell cycle and cellular senescence) and four signaling pathways (AMPK, JAK-STAT, p53, and PI3K-AKT). Using Fisher’s exact test between the genes in a pathway and a TF’s regulon (here, we consider the TF and its targets), we compute significance of overlapping between them and annotate each TF in the optimal network with the most significant pathway (**Fig. 4c**). The coupling between these pathways is shown in **Fig. S2.** JAK/STAT is the central communication node in cell function that is involved in cellular progression and differentiation together with hematopoiesis among other functions^41^. In a recent study, Habbel et al. found that JAK/STAT signaling pathway is activated because of the inflammation in the AML cells^42^. Also, AML enables the myeloid cells to proceed uncontrolled and limitless number of cell cycles^43^. Cellular senescence promotes the evasion of tumor cells from immunosurveillance^44^ . The coupling of JAK-STAT signaling pathway and cell cycle suggests increased cell-cell communication and expedited cell growth, which is shown in recent in vitro experiments^45^. On the other hand, the activation of p53 signaling pathway coupled with cellular senescence can be attributed to the DNA damage and subsequent cell cycle arrest in leukemogenesis ^46, 47^. PI3K-AKT signaling pathway is found to play a role in both cell proliferation^48^ and cell cycle arrest^49^ in AML. AMPK exhibits a dual role in AML, as it acts as a tumor suppressor before the disease onset but can promote disease progression after its onset in association with other key pathways^50^. Together, the findings suggest that the coupled gene regulation of these signaling pathways contributes to tumorigenesis in AML.

### Perturbation analysis reveals significant TFs in the optimal GRN

With the established optimal GRN, simulations of gene perturbations can be performed to identify crucial TFs or TF pairs destabilizing the network states^16, 51, 52^. Here, we simulated the GRN with either single or double gene knockdown (KD), and, for each case, we evaluated the proportion of models belonging to the normal and the AML states of the GRN (**Supplementary Note 6**). When the proportion of models in the AML state increases, the gene(s) undergoing KD would be regarded as destabilizer(s) of the AML state. From single KD perturbations, the top five destabilizers of the AML state are TFDP1, E2F4, TP53, MYC, and E2F1; in contrast, the top five destabilizers of the normal state are STAT3, RB1, POU2F1, ETS2, and MYCN. These top 10 destabilizers are associated with three key biological pathways: JAK-STAT signaling (STAT3, POU2F1), Cell cycle (TFDP1, E2F4, MYC, E2F1, RB1, ETS2, MYCN), and p53 signaling (TP53). Activation of JAK-STAT signaling and cell cycle indicates increased cell cycle communication and cell growth^45^, requiring activation of p53 signaling for repairment of increased DNA damage^46^. These top destabilizers from both directions were then used for double KD simulations. As expected, the double KDs have higher impact to the network states than the single KDs (**Fig. 6a**). Among all of the single and double KD simulations, 10 double KD perturbations were found to significantly expand the model proportions of the AML state (by a Chi-squared test, lower part of **Fig. 6b**).

**Figure 6.**
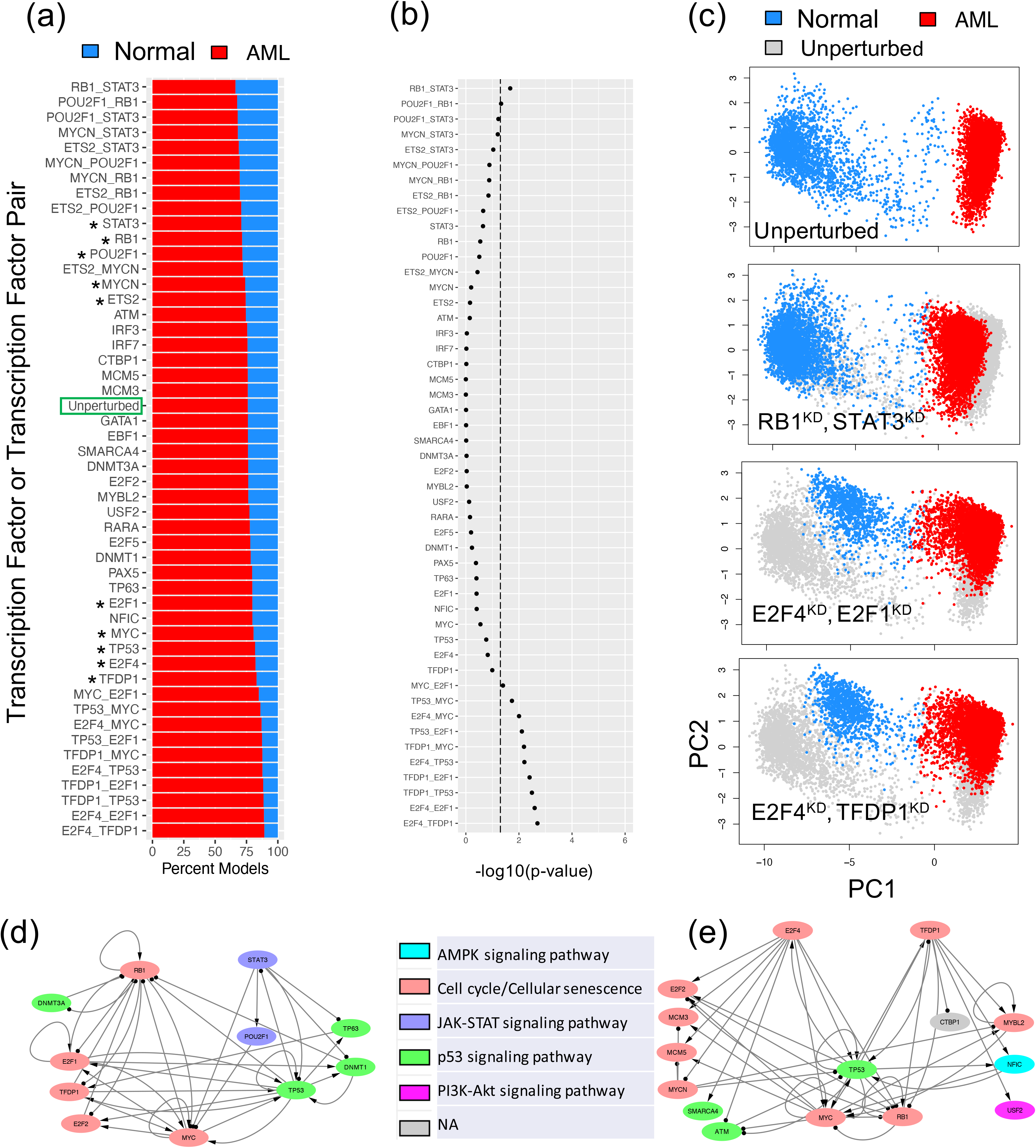
In silico perturbation analysis of the AML GRN. **(a)** Proportion of models (belonging to the two experimental groups -- normal controls and AML patients) from single and double knockdown (KD) simulations. Perturbations are arranged in descending order based on model proportions in the normal group. The top five and bottom five genes (marked by stars) were used for double KD simulations. **(b)** Significance of changes in gene expression states upon GRN perturbations by a chi-squared test. X-axis represents −log10(p-value). Dotted line indicates p-value=0.05. **(c)** Examples of changes in gene expression profiles upon GRN perturbations. The first row shows the scatter plot of the simulated gene expression profiles of the GRN under the unperturbed condition projected on the first two principal components of the data. The second to the fourth rows show the scatter plots of the simulated gene expression profiles of the GRN under various perturbed conditions in the same PCA space. For the last three rows, perturbed expressions are shown (blue: normal, red: AML) on top of the unperturbed expressions (gray). Top single KD perturbations are shown in **Fig. S3**. **(d)** A subnetwork containing RB1, STAT3, and their target transcription factors. A double KD of RB1 and STAT3 causes the largest decrease of the models in the AML state. **(e)** A subnetwork containing E2F4, TFDP1, and their target transcription factors. A double KD of E2F4 and TFDP1 causes the largest increase of the models in the AML state.

Furthermore, we examined in detail how the network states change for the top three KD perturbations (*i.e.*, RB1-STAT3; E2F4-E2F1; E2F4-TFDP1) (**Fig. 6c**). First, we performed principal component analysis of the RACIPE-simulated gene expression profiles for the unperturbed condition and projected those profiles onto the first two principal components (PCs) (top panel in **Fig. 6c**). Next, the KD simulated gene expression profiles were projected on the same PCs, as shown in the bottom three panels in **Fig. 6c** and **Fig. S3**. Noticeably, the double KD of the TF pair RB1-STAT3 shifts the gene expressions of the AML models towards those of the normal models. On the other hand, the other two double KD perturbations, E2F4-E2F1 and E2F4-TFDP1, shift the gene expressions of the normal models towards those of the AML state. Hence, the perturbation analysis of the optimal GRN reveals the significant TFs and TF pairs that can shift the cell populations from AML state to normal state and vice versa. Such information can be important in designing effective therapeutic strategies.

To further examine the synergistic effects of the TF pairs in the double KD perturbations, we checked the two subnetworks consisting of the targets of TF pair RB1-STAT3 and TF pair E2F4-TFDP1, as shown in **Figs. 6de**. Here, the double KD of RB1-STAT3 has the largest impact to destabilize the AML state, while the double KD of E2F4-TFDP1 has the largest impact to destabilize the normal state. The E2F4-TFDP1 KD causes larger changes possibly because both TFs are on the same pathway and have a higher number of overlapping target nodes, MYC, RB1, and TP5, in the GRN (**Fig. 6e**), whereas only one overlapping target node MYC for RB1 and STAT3 (**Fig. 6d**).

### Survival analysis suggests therapeutic strategies

To investigate the relationship of the 29 TFs in the GRN with the prognosis of AML patients, we performed Kaplan-Meier survival analysis and log-rank test. We performed the survival analysis for two scenarios: in one case, we used only nine IDH mutant AML patients and, in the other case, we used all 119 AML patients. In each case, we calculated the risk score for each patient using the expression profiles of each individual TF and its target genes. We divided the AML patients into two groups (high risk and low risk) based on their risk scores. For the key TFs, such as E2F1, NFIC, and TP53, a significant difference in event-free survival was observed between high- and low-risk groups (**Figs. 7, S3**). Additionally, these TFs were also found to be among the most impactful genes in the KD simulations (**Figs. 6abc, S2**). These results suggest that the identified TFs could act as prognostic factors of leukemia. Our observations are also supported by existing literature on AML studies. Pulikkan et al. showed that E2F1 forms an autoregulatory negative feedback with miR-223, and inhibition of miR-223 increases myeloid cells in AML^53^. Thus, overexpression of E2F1 can increase AML severity. In another recent study, Dutta et al. analyzed the TP53 mutation profiles of AML patients and found that AML patients with TP53 mutations showed worse prognosis than patients with wild type TP53^54^. GATA1, another prognostic factor found in our analysis, was also reported to be overexpressed in AML^55^. This analysis further supports that the optimal GRN included important TFs that are not only significant for IDH1/2 mutant AML leukemogenesis, but also predictive for the survival of other types of AML patients.

**Figure 7.**
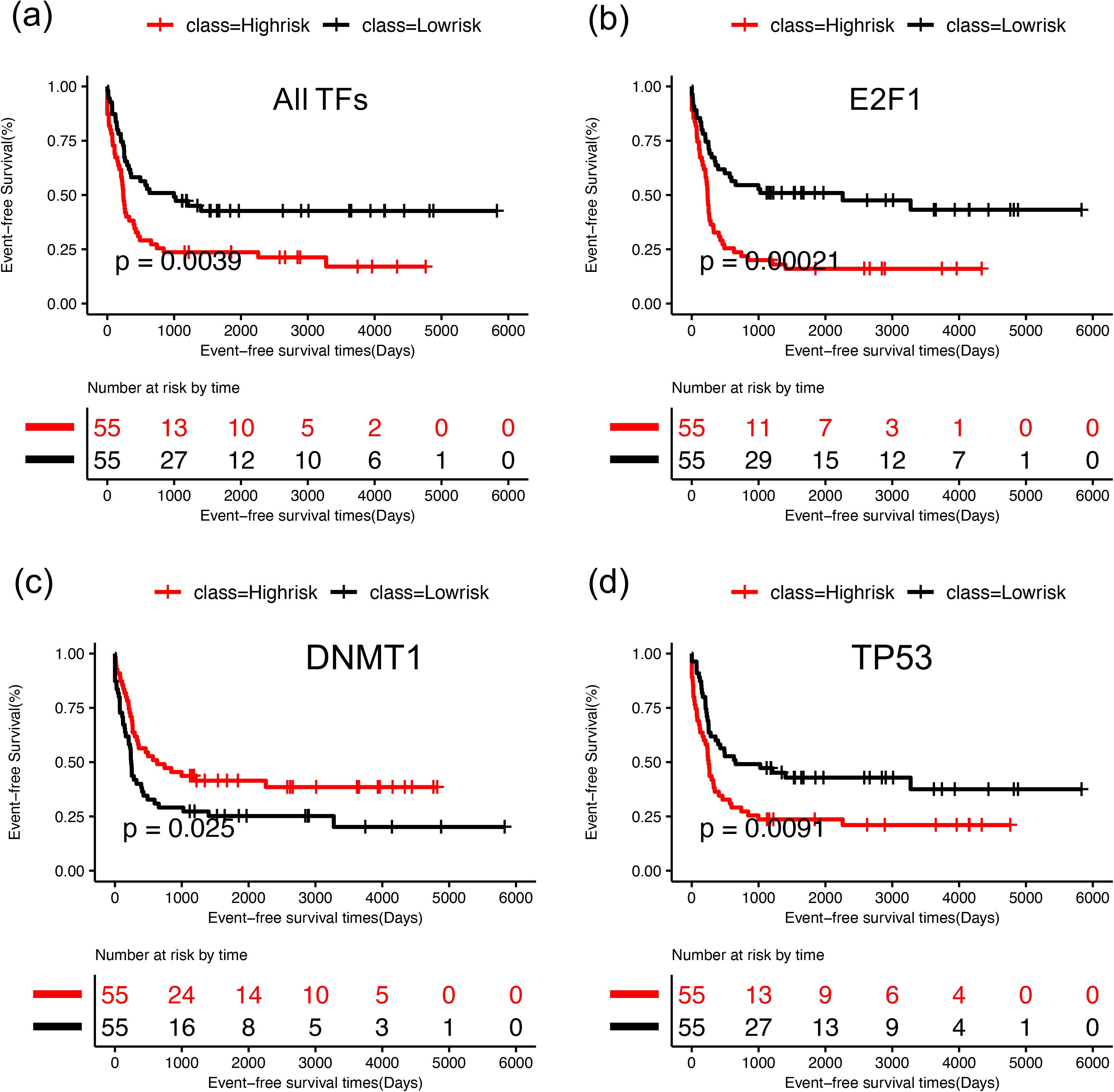
Survival Analysis of the optimal GRN. For each TF, log-rank test was performed to group the patients as high (red) and low risk (black) based on median of risk scores and then Kaplan-Meier analysis was performed for survival analysis. **(a)** Kaplan-Meier curves for event free survival for E2F1 using nine IDH AML patients, **(b)** Kaplan-Meier curves of event free survival for E2F1 using 119 AML patients, **(c)** Kaplan-Meier curves for event free survival for NFIC using 119 AML patients, **(d)** Kaplan-Meier curves of event free survival for TP53 using 119 AML patients. See **Fig. S4** for survival analysis for additional TFs using all 119 AML patients.

## Discussion

With the advent of high-throughput sequencing technology, large datasets of transcriptomic, proteomic, and genomic profiles of cancer patients, together with literature-curated gene regulatory interactions, have been available. Identifying the differentially expressed genes for cancer subtypes and the related enriched pathways does not clearly inform us the underlying gene regulatory mechanism of molecular state change in tumorigenesis. Despite the availability of plethora of molecular profiles of tumor samples, there is still a lack of suitable methodologies to extract important information from the diverse tumor datasets for a mechanistic understanding of tumorigenesis. Several top-down bioinformatics methods utilized high-throughput gene expression data to study dysregulation of gene expression in cancer^56–58^ and link the upstream signaling pathway to downstream transcription program^59^. Some other methods infer network of transcription factor and target genes^31, 60, 61^. Although the regulatory maps inferred by these methods give a global view of gene regulation, the generated networks usually do not capture the gene regulation of the state transition between normal and cancer cells^14^. To address this issue, there is a need to develop approaches that allow to establish systems-biology gene network models for predicting gene expression dynamics directly from diverse cancer genomics data sets.

Here, we introduced a generic computational framework by extending our recently published method, NetAct^62^ for modeling GRNs driving cellular state transitions during disease development by using a combined top-down bioinformatics and bottom-up mathematical modeling approach. The top-down approach was applied to generate a collection of putative GRNs by integrating genomics data from diverse sources. Subsequently, the bottom-up mathematical modeling approach was applied to identify the optimal GRN that reproduces experimental gene expression data. Compared to NetAct, the method presented here offers two key enhancements. First, it integrates ATAC-seq data and literature-based curated TF-to-target gene relationships, whereas NetAct solely relies on the curated database. Second, the current method employs mathematical modeling to identify the optimal gene regulatory network (GRN) among many candidate GRNs. Empowered by these improvements, the current method enables us to find the optimal GRN that elucidates the gene regulatory mechanism of leukemogenesis in AML and unravels the coupling of relevant biological pathways. In particular, the method successfully captures a key regulator DNMT1, a known factor associated with IDH1/2 functions^35^. The optimal GRN also identifies key genes involved in cell cycle regulation and DNA damage repair, such as RB1, E2F1/2, TP53, and MYC, along with stem cell pluripotency factors GATA1, POU2F1, and MYCN. Overexpression of these genes suggests that AML cells acquire a stem cell-like phenotype with a restricted cell cycle, potentially leading to drug resistance. In addition, the single and double knockdown simulations of the GRN identified E2F1 as one of the top TFs whose knockdown significantly increased the cancer state, which is supported by the survival analysis of the AML patients.

While our approach has yielded promising results, several limitations warrant investigation for future advancements. We currently applied our approach to study AML tumorigenesis whereas the dataset captures mainly two cellular states. It would be interesting to apply such an approach to systems where one or multiple intermediate states are captured in the data. Additionally, the integration of multiomics datasets, such as microarray gene expression data and ATAC-seq chromatin accessibility data obtained from separate experiments, may benefit from the generation of multimodal datasets, where both datasets are obtained from the same cells. Such integration would enhance the context-specificity of inferred GRNs. Furthermore, other valuable data types, like Hi-C data, could offer regulatory information not currently accounted for in our method. Another consideration pertains to the time-consuming nature of simulating all potential GRNs to identify the optimal network, especially when dealing with a substantial number of inferred GRNs. This can be mitigated by parallelizing the simulations of potential GRNs, which can significantly reduce the computation time. Implementing this parallelization would enhance the efficiency and scalability of our approach, making it more practical for larger datasets and complex analyses.

Despite these limitations, our current approach marks a valuable steppingstone in exploring gene regulatory networks as systems biology network models. Addressing these considerations in future research will undoubtedly improve the method’s capabilities, enabling it to deliver even more comprehensive and accurate insights into the regulatory mechanisms of cellular state transitions.

## Authors’ Disclosures

No disclosures were reported by the authors.

## Authors’ Contributions

**A. Katebi:** Formal analysis, investigation, methodology, writing with inputs from others. **X. Chen:** Formal analysis, investigation, methodology, writing. **S. Li:** Conceptualization, supervision, funding acquisition, writing – review and editing. **M. Lu:** Conceptualization, supervision, funding acquisition, writing – review and editing.

## Acknowledgments

The study is supported by startup funds from The Jackson Laboratory and Northeastern University, by the National Cancer Institute of the National Institutes of Health under Award Number P30CA034196, and by the National Institute of General Medical Sciences of the National Institutes of Health under Award Number R35GM128717.

## Notes

### Competing Interest Statement

The authors have declared no competing interest.

