## Supplemental Materials for "Data-driven modeling of core gene regulatory network underlying leukemogenesis in IDH mutant AML"

### Supplemental Information

#### Supplementary Note 1. Selecting TFs using three distinct bioinformatics methods

A list of TFs was obtained by applying each of the three previously published methods: Virtual Inference of Protein-activity by Enriched Regulon analysis (VIPER) [2], Regulatory Inference (RI) [3], and NetAct [4].

**Preprocessing Rcistarget data for VIPER and RI methods**. The cis-binding motifs for human transcription factors were collected from Rcistarget v1.3, which contains 982 transcription factors (TFs) and 1872 motifs. Position weight matrices (PWMs) were converted to the MEME motif format[1] and the FIMO tool from the MEME package was used to search for binding sites at the open chromatin peaks within 2 kb upstream and downstream of the transcription start sites (p-value<0.0005). We used the default parameter of FIMO except for the max-stored-scores and motif-pseudo-options which we set to 100,000,000 and $1\times10^{-8}$, respectively.

**VIPER:** First, we used the function *aracne2regulon* from ARACNe algorithm[5] to generate context-specific regulatory network based on gene expression of AML patients with IDH mutation and $CD34^+$ controls. Then, the *msviper* function in viper R package is used to generate normalized enrichment score (NES) and p-value, which identified 230 key IDH-specific TFs with FDR-adjusted p-value less than 0.05.

**RI (sample-by-sample lasso regression models):** We used sample-by-sample lasso regression models from the RI method [3] with inputs gene expression profile and regulatory sequence information to infer sample-specific TF activities and IDH-specific key regulators. Here, we used linear regression to model log gene expression changes in AML patients with IDH mutation versus $CD34^+$ controls by TF binding site counts in the gene promoter as variate. Quantification of binding site counts from ATAC-seq data can be found in the method section named *ATAC-seq data*. Lasso regression was performed using the glmnet function in the R package[6]. The regularization parameter was determined using 10-fold cross validation for each sample. The coefficient of each TF estimates the importance of the TF in the sample. We performed feature dependency analysis using RI method to obtain 938 key IDH-specific TFs.

**NetAct:** We employed our newly developed method NetAct[7] for TF selection from the gene expression data (11 normal controls and 9 AML patients with IDH mutations, GEO accession number GSE6891)[8,9] and TF-target gene database. First, a two-way comparison (normal control and IDH mutation condition) was performed for differential gene expression (DE) analysis using limma[10] (in NetAct, we provide function *DEG_Analysis_Micro*). This generated a ranked gene list quantified by adjusted p-value. Then, the enriched TFs were identified by performing gene set enrichment analysis (NetAct function TF_Selection with slight modification on GSEA, number of permutations=1000) using our curated TF-target gene database. The curated TF-target gene database was compiled from different sources: TRRUST[11], RegNet[12], and TFactS[13], TRED[14], FANTOM5[15], ChEA[16], TRANSFAC[17], JASPAER[18], ENCODE[19], and RcisTarget[20]. For GSEA, we considered 312 TFs with eight or more targets in the TF-target gene database[7] and obtained the TFs ranked by adjusted p-value.

**Supplementary Note 2. Combining literature-based TF-targe genes and ATAC-seq data**

We constructed TF-target databases by combining curated TF-target gene database with TF-target gene relationships obtained from the ATAC-seq dataset at different TF-target binding probability thresholds, with the aim of finding a balanced mix of curated and ATAC-seq targets. At each TF-target gene binding probability threshold, we selected the targets for each TF from the ATAC-seq data according to the following criteria:

$$n_{target} = \begin{cases} n_{genes}, \; n_{genes} < N_{TSH} \\ N_{TSH} + (n_{genes} - N_{TSH}) * p, \; n_{genes} > N_{TSH} \end{cases}$$

where $n_{genes}$ represents the number of probable target genes above the TF binding probability threshold, $N_{TSH}$ represents the threshold for the number of probable target genes below which all are selected as targets (set at 50), and p represents the percent of genes used to select the top target genes from the probable target genes ($n_{genes}$) (set at 0.01 to select the top 1% target genes). Then, the inferred TF-target gene relationships are a specific TF-target gene binding probability threshold were merged with the curated TF-target gene database. We retained the TFs with at least eight targets in the merged TF-target gene database. Eleven TF-target gene binding probability thresholds were chosen: 0.05, 0.06, 0.07, 0.08, 0.09, 0.10, 0.12, 0.14, 0.16, 0.18, and 0.20 (**Table S1**).

**Supplementary Note 3. Constructing candidate GRNs and identifying the optimal GRN**

To construct candidate GRNs, we first inferred a list of core TFs as follows. First, we selected a specific number of TFs from each of the three bioinformatics methods (NetAct, VIPER, and RI). We ensured that each selected TF has at least eight target genes in the merged TF-target gene database. Then, we combined the TFs selected from each method at each ATAC-seq TF-binding probability cutoff. See **Table S1** for the choice of the hyperparameters number of TFs per method and ATAC-seq TF-binding probability cutoff. The combined set of core TFs selected at each ATAC-seq probability cutoff and TF count per method, were then used for putative GRN construction as described below.

From a set of core TFs, we constructed an initial network set by connecting any TF pair from the combined TF-target gene database. Each regulatory interaction contains a regulator TF and target TF, and the interaction type could be either excitatory or inhibitory, determined by the sign of the Spearman correlation between the activities of the regulator and target TF pair. Only those interactions are retained whose absolute correlations are above a given threshold value. If the obtained network consists of multiple disconnected subnetworks, we retained the largest subnetwork containing more than 80% of the TFs from the obtained network. If the largest subnetwork is smaller than 80% of the obtained network, we discarded the network. We repeated the above process for 20 TF activity Spearman correlation cutoff starting from 0.0 to 0.95 with a 0.05 stepwise increment. In this way, we retained 532 candidate GRNs with 15 or more TFs per network.

We obtained the optimal network among the 532 candidate GRNs according to the combined accuracy and flexibility ranking (as defined below in **Supplementary Note 4**). For each candidate GRN, 10000 RACIPE models were first generated to compute the accuracy and flexibility. The candidate GRNs were then ordered by the accuracy and flexibility (both from high to low), respectively. The combined index of a GRN was defined by the sum of the ordering indices of the accuracy and flexibility.

**Supplementary Note 4. Accuracy and flexibility**

Two metrics, accuracy and flexibility, were used to rank the candidate GRNs in the network optimization process. Accuracy captures the context specificity of a GRN by matching the RACIPE simulated gene expression with the experimental gene expression data, whereas flexibility captures the plasticity of the network by contrasting the RACIPE simulations under the unperturbed and perturbed conditions.

Accuracy of a candidate GRN was measured by the fraction of the RACIPE models (under the unperturbed condition) that can be assigned to any of the two experimental gene expression states (normal controls and AML patients). To assign a RACIPE model to an experimental state, first we calculated the Euclidean distance between the simulated gene expression profile of the RACIPE model and the nearest TF activity profile of a sample from the experimental state. Second, we generated 1000 random gene expression profiles by shuffling gene names and calculated each profile's distance to the nearest TF activity profile of a sample from the experimental state. Using the distances from the random profiles as the null distribution, we calculated the p-value for each RACIPE model to be in that experimental state. Finally, we mapped each RACIPE model to the experimental state with the smallest p-value. If the p-values corresponding to all experimental states are greater than 0.05, we considered the RACIPE model to be unassigned, indicating that the model could not be mapped to any experimental state.

Flexibility of a GRN was defined as the differences in the distribution of the assigned gene expression states of an ensemble of 10000 RACIPE models between the unperturbed condition and any single-gene knockdown (KD) condition. The formula to compute the flexibility is

$$flexibility = \frac{1}{n}\sum_{i=1}^{n} sqrt\left(\left(m_{normal}^{u} - m_{normal}^{KD_i}\right)^2 + \left(m_{AML}^{u} - m_{AML}^{KD_i}\right)^2\right)$$

where n is the total number of TFs in the candidate GRN, $m_{normal}^{u}$ ( $m_{AML}^{u}$) is the proportion of the RACIPE models mapped to the normal control (or AML) experimental state under the unperturbed condition, and $m_{normal}^{KD_i}$ ( $m_{AML}^{KD_i}$) is the proportion of the RACIPE models mapped to the normal control (or AML) experimental state under the KD condition of the $i^{th}$ TF.

**Supplementary Note 5. Annotating the TFs in the optimal GRN with significant biological pathways**

We annotated the most representative biological pathways to the GRN TFs as follows. First, we obtained the differentially expressed genes (DEGs with adjusted p-value <0.05) between the two groups normal controls and AML patients and only retained the DEGs that were either TFs in the network or their targets found in the corresponding TF-target gene database. Second, we applied enrichr [21] to find 12 enriched KEGG pathways from the DEGs (adjusted p-value < 1.60e-10). From these 12 pathways, we disregarded five pathways (Pathways in cancer, Epstein-Barr virus infection, Hepatitis B, Measles, and Human papillomavirus infection), because they were either too generic (Pathways in cancer) or not directly related to AML (Epstein-Barr virus infection, Hepatitis B, Measles, and Human papillomavirus infection). We selected the following seven top-ranked pathways from the enrichment analysis: Cell cycle, p53 signaling pathway, PI3K-Akt signaling pathway, JAK-STAT signaling pathway, MAPK signaling pathway, Cellular senescence, and AMPK signaling pathway. We performed Fisher's exact test to check whether genes from each pathway overlaps with the DEGs corresponding to each TF (both the TF and their targets in the TF-target gene database) (**Fig. 1b**, **Table S2)**. Finally, we annotated each TF with the pathway that has the smallest p-value provided that the p-value ≤ 0.1. If no significant pathway with that p-value threshold was found for a TF, the TF was unassigned to a pathway (**Table S2**). The annotated GRN was then visualized using Cytoscape [22], as shown in **Fig. 4c**.

**Supplementary Note 6. Characterizing the dynamics of optimal GRN**

Single and double knockdown simulations were performed on the AML GRN using RACIPE method[23–25] as follows. First, 10000 RACIPE models were simulated for the AML GRN to generate the gene expression profiles for the unperturbed condition. Second, for the single KD simulations for a TF in the network, we reduced the production rate of the corresponding TF by 95% for each RACIPE model and then re-simulated the model to generate the gene expression profiles for the KD condition. Third, for double KD simulations, for each RACIPE model, we reduced the production rate of both TFs by 95% and then re-simulated the model to generate the gene expression profiles for the double KD condition. We used ridge regression to map the knockdown RACIPE simulated expressions to the two groups, normal controls and AML patients. To achieve this, first, we mapped the 10000 RACIPE models from the unperturbed simulations to normal controls and AML patients using the method described in **Supplementary Note 4** and used these labeled unperturbed models to train a regression model. We then used the trained regression model to map the knockdown RACIPE simulated expressions. Afterwards, we calculated the proportion of RACIPE models mapped to each group, normal control and AML patient. The effect of TF knockdown was evaluated by the change in the proportion of the models matching the two experimental groups normal controls and AML patients, compared to the simulations from the unperturbed condition.

### Supplemental Tables

**Table S1. The choice of optimization hyperparameters**

| **Parameters** | **Values** |
|---|---|
| TF binding probability (ATAC-seq) | 0.05, 0.06, 0.07, 0.08, 0.09, 0.10, 0.12, 0.14, 0.16, 0.18, and 0.20 (11 values) |
| Number of TFs per method (NetAct, VIPER, RI) | 4 to 60 at an interval of 4 (15 values) |
| TF activity correlation cutoff | 0.0 to 0.95 at an interval of 0.05 (20 values) |

**Table S2. Overlapping between TF regulons and top enriched biological pathways**
First column shows the gene symbol for each TF in the optimal GRN. Second column shows the names of top enriched pathways and the p-values for the Fisher's exact test between a TF regulon (TF and its target genes) and genes in a pathway.

(See Table-Se.TFsWithAnnoted_pathway.xlsx)

### Supplementary Figures

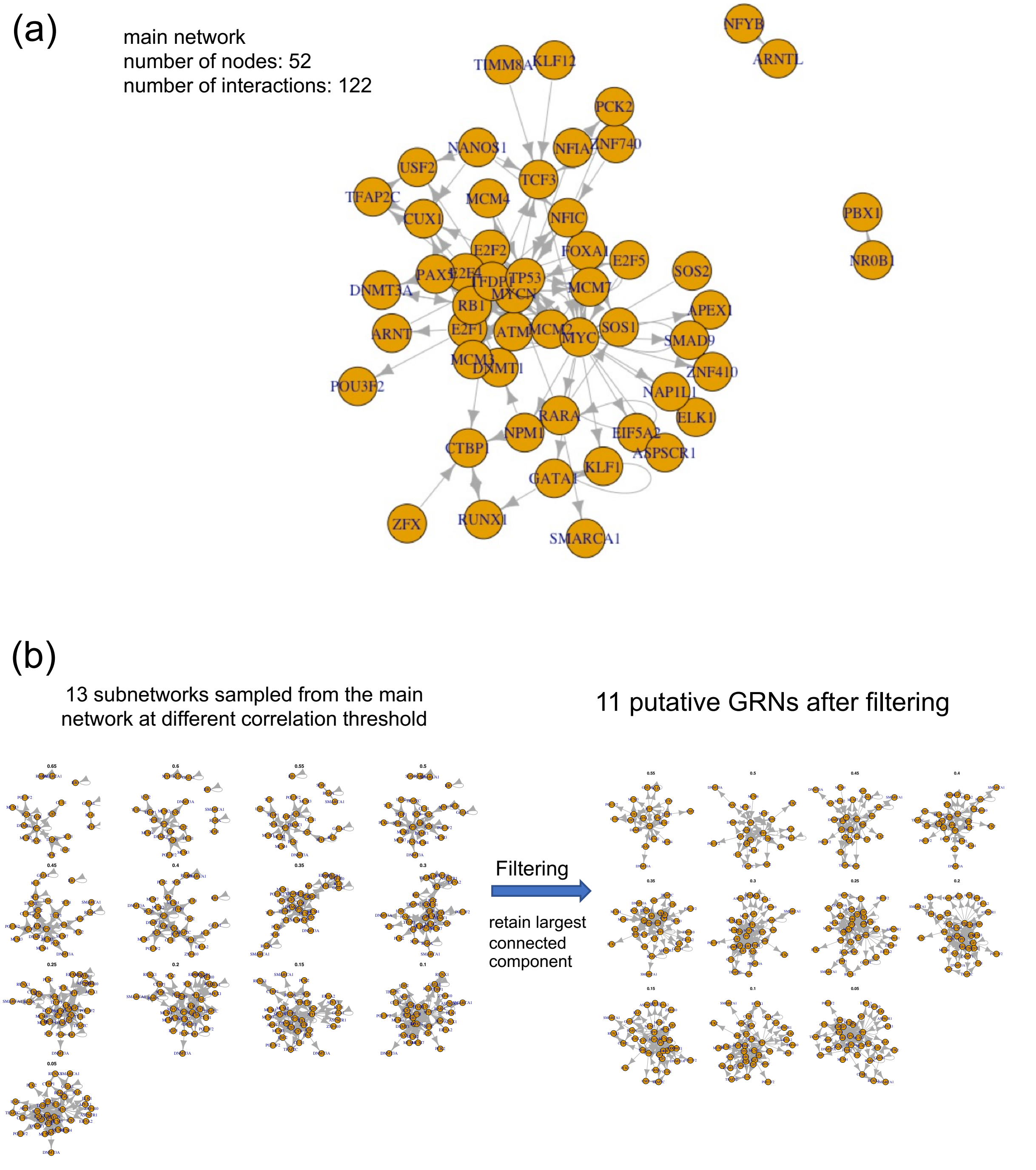


**Fig. S1 (related to Fig. 1). Inferred putative GRNs. (a)** Initial network with 52 nodes and 122 interactions. **(b)** Left plot shows 13 GRNs obtained at different TF activity correlation threshold from the initial network shown on panel **a**. Right plot shows, from the 13 GRNs, the largest subnetworks, each of which have more than 80 percent of the nodes of the corresponding sampled network.

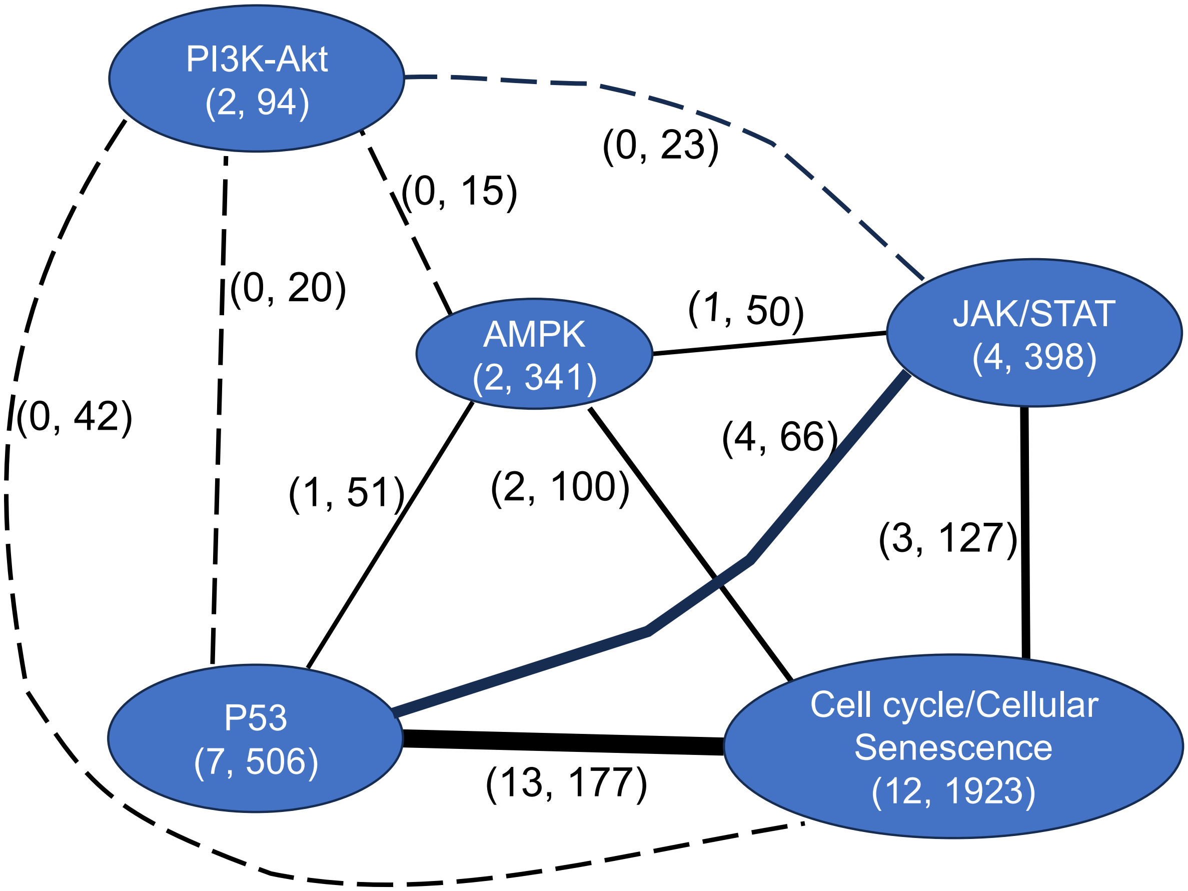


**Fig. S2 (related to Fig. 4).** Coupling of biological pathways. Nodes represent pathways as labeled. The two numbers within each node represent the number of transcription factors (TFs) corresponding to that pathway and the total number of genes targeted by those TFs, respectively. The two numbers on each edge label represent number of links between the two TF groups and the number of common target genes targeted by the TF groups. Thicker edges indicate comparatively more TF links between the two groups. Dotted edge indicates no links found between the two TF groups.

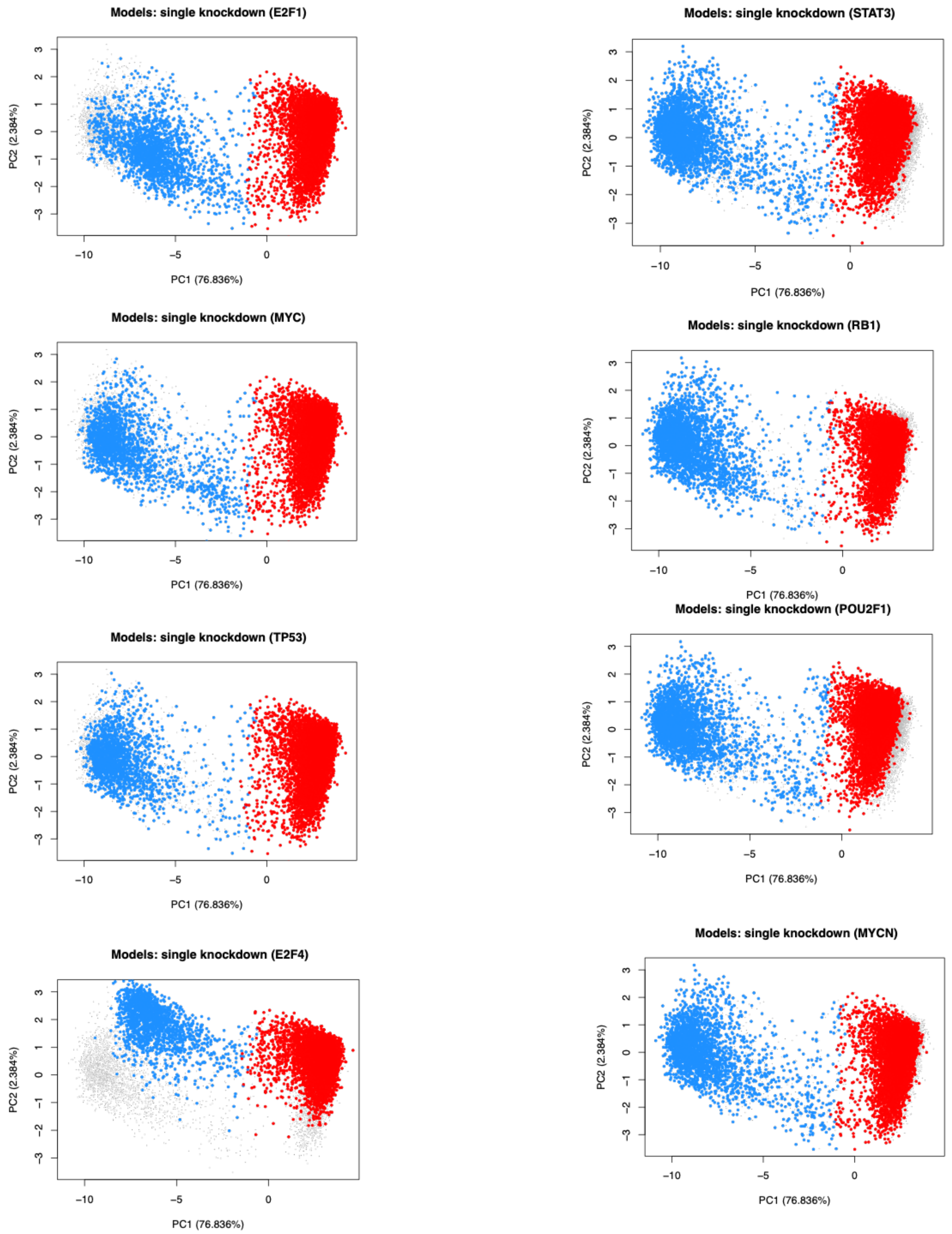


**Fig S3 (related to Fig. 6c)**. Examples of changes in gene expression profiles upon single knockdown perturbations. Left: Perturbations that increase AML proportions. Right: Perturbations that decrease AML proportions.

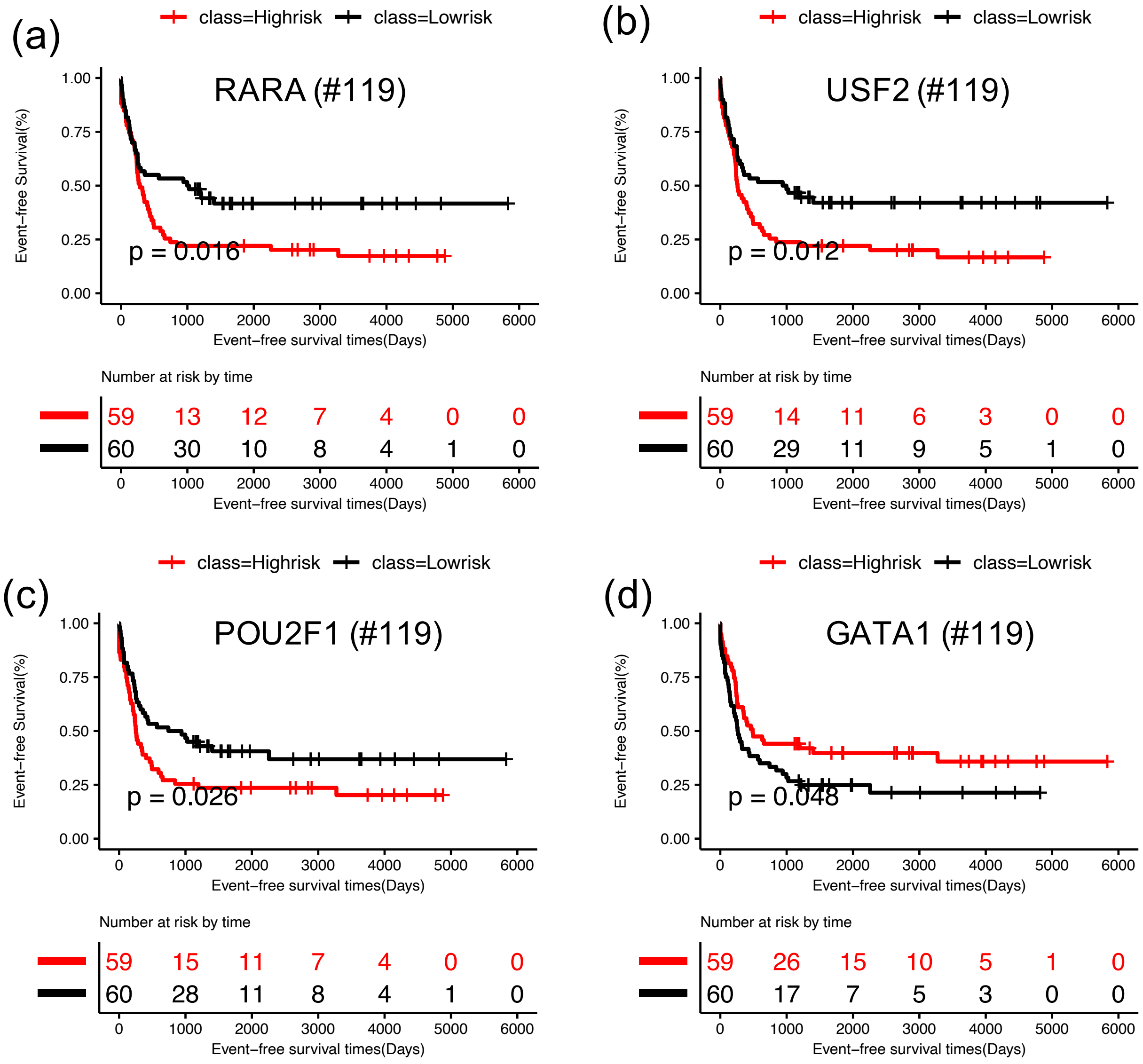


**Fig. S4 (related to Fig. 7).** Kaplan-Meier curves for event free survival for individual TFs on the AML GRN using all 119 AML patients with p-value <= 0.05. (a) RARA, (b) USF2, (c) POU2F1, and (d) GATA1.
